# Repurposing of Drugs Against Mutated Strain of Eurasian Avian Like H1N1 (EA H1N1) Swine Flu Virus, Genotype 4(G4) Virus

**DOI:** 10.1101/2022.10.20.512704

**Authors:** Sangita Ghimire, Sandhya Sahukhal, Ayush Shrestha, Sarmila Adhikari, Samiran Subedi, Keshab Raj Budha, Pramod Aryal

## Abstract

Mutation, reassortment and recombination have led to the evolution and the emergence of more pathogenic and new subtypes of influenza virus. The surge of highly mutated viruses has prompted the need of coherent solution for the so called “medical holocaust” viral outbreaks. The genotype 4 of EAH1N1 strain has been circulating in the swine population as a dominant genotype, exhibiting even human to human transmission. This has risen the possibility of causing another global health threat as a lethal viral outbreak in the future. The Computer Aided Drug Discovery (CADD) could be a prudent mechanism to develop new drug candidates against such disease for its mitigation. In this regard, the computational *in silico* methods had been envisaged in this research for the prediction of lead compounds against the selected proteins of EA H1N1 G4 strain, namely Haemagglutinin (HA) and Polymerase acidic protein(PA). The research focused on the selection of the target viral protein and molecular docking for the identification of putative ligands. It was followed by the identification of the probable mutations and assessment of effectiveness of identified drugs against their respective targets. Total of 3 compounds Enalapril, Enalaprilat and Ivabradine have been identified as a potential inhibitor of HA and PA protein that were prioritized on the basis of preference index parameter and binding energy of compound with the respective target. Besides, the probable mutations in each target protein in future were predicted and all these 3 top hits were found to be effective against mutated variant of these proteins. Thus, Enalapril, Enalaprilat and Ivabradine could be the lead compounds to explore further as multi target inhibiting drugs against wild and mutant variant of target proteins.

## 1. Introduction

The influenza virus is an enveloped, 8 negative sense single stranded RNA influenza virus of family *Orthomyxoviridae* and genus Influenza virus that comprises common human flu(Nogales et al. 2018), bird flu, swine flu(Sahoo et al. 2016) and others (Webster and Govorkova 2014)responsible for acute respiratory disease (Abbas and Abidin 2013, Vasin et al. 2014). The influenza A virus has been classified into 16 HA subtypes till date (Li et al., 2011). The H1N1 is a subtype of Influenza A virus that also is in the group of swine flu virus responsible for deadly influenza pandemic in the past (Chen et al. 2009).

The emerging global threat of influenza viruses is the result of continuous evolution due to various reasons. Mutation (antigenic drift), reassortment (antigenic shift) and recombination are the major mechanism responsible for the evolution in the influenza virus (Shao et al. 2017).The formation of new subtypes of influenza virus plays a significant role in wide propagation and disastrous outbreak as there is dearth of immunity to the newly emerging subtypes (Ma, Kahn, and Richt 2008). The vaccination has been used as a staple method against epidemic and pandemic influenza. However it is not able to provide the immediate protection against such epidemic and pandemic flu outbreaks. Thus, it is prudent to find the treatment to contain the lethal pathogenic viruses outbreak to prevent a major global health threat.

In this aspect, an influenza virus surveillance of pigs from 2011 to 2018 in China (Sun et al. 2020)revealed a newly emerged genotype 4 (G4) Reassortant Eurasian avian-like (EA) H1N1 virus with 2009 pandemic (*pdm/09*) and triple-Reassortant (TR)-derived internal genes (Sun et al. 2020) as predominant in swine population since 2016 and that the EA H1N1 with highest potential to cause the next lethal outbreak (Yang et al., 2016) demands strict monitoring. A predominant emergent of G4 swine flu virus has the glycoprotein Hemagglutinin (HA) and Neuraminidase (NA) originated from EAH1N1 and other proteins Matrix protein (M), Nucleoprotein (NP), Nonstructural protein (NS), Polymerase basic protein 1 (PB1), Polymerase basic protein 2 (PB2).

The antigenicity of the two cell surface glycoprotein HA and NA have to be taken in consideration as these are the basis of subtype classification of G4 EAH1N1 (Hoffmann and Pöhlmann 2018) and HA is one among the two surface envelope glycoproteins responsible for attachment of the virions to the host receptors as well as the role in releasing newly produced viral particles. The swine viruses possess ability to bind sialic acid in both α-2,3 and α-2,6 linkages (Gamblin & Skehel, 2010) having properties of avian/equine influenza viruses that preferentially binds to the sialic acid in α-2,3-linkage to galactose (Ito et al. 1998)and human influenza virus that preferentially bound to α-2,6-linked sialic acid(Ito et al. 1998). The initial process of infection occurs when the globular head of the HA binds to the sialic acid receptors present on the host membrane and virion and the lower pH facilitates the cleavage of the cylindrical tail from the globular head for host endosome membrane and viral membrane to fuses together releasing viral RNA and protein into the cytoplasm of the host (K. P. Le et al., 2019; Russell et al., 2018) to internalize through endocytosis. Thus, G4 EAH1N1 influenza virus protein, HA is a good target for treatment and prevention of transmission.

Similarly, polymerase acidic protein (PA) from A(H1N1) pdm09 (Sun et al. 2020) is also of concern because of the reassortments and requires advance preparedness with possible treatment. PA is phosphoprotein that functions as endonuclease and bound by PB2 is cleaved downstream of their mRNA cap structure (S. Yuan et al. 2015). It also acts as protease due to the presence of a third amino terminal nearer to its nuclear localization signals (Abbas and Abidin 2013). PA is divisible into smaller N-terminal (25k Da) domain and larger C-terminal (55k Da) by trypsinization (P. Yuan et al. 2009). The N-terminus of the PA subunit consists of the endonuclease domain. (Kowalinski et al. 2012). PA mutation also shows to be connected to the adaptation of porcine H1N1 virus to new host species (Bhoye and Cherian 2018).

Thus, identification of medical entity that could be used as antviral agent(s) would be vital to tame the outbreak, if any that may arise. For rapid identification of medical entity computer aided drug discovery(CADD) could be an efficient means of identifying potential lead compound as CADD is suggested for development of possible drugs for a wide range of diseases and accelerate the process of drugs discovery (Baig et al. 2017). In addition, drug repurposing, a strategy for identifying new uses for approved or investigational drugs could be employed (Kumar et al. 2019).

## 2. Methods

### 2.1 Selection of target viral protein and retrieving their genomic sequences

HA protein was chosen as a drug target due to its role in binding of virus to the upper respiratory tract of human, mediating viral entry and fusion(Le et al., 2019).

Polymerase Acidic (PA) protein was selected as target protein as it is RNA dependent RNA polymerase with several roles such as endonuclease function, protease activity and viral RNA/ complementary RNA promoter binding (Yuan et al. 2009).

The genomic and protein FASTA sequences of haemagglutinin and polymeric acidic protein of EAH1N1 G4 virus were retrieved from the supplementary materials of the research(Brázda et al., 2021)

### 2.2 Homology modeling and protein preparation

The tertiary structure of HA and PA protein were modelled by homology modeling on the basis of the available template that had been deposited in the Protein Data Bank (PDB)(https://www.rcsb.org/)(Berman et al., 2000).The template 6D8W(2.35 Å) (https://www.rcsb.org/structure/6D8W was used as a homolog template to build a HA model using Swiss model(https://swissmodel.expasy.org/)(Schwede et al., 2003)Similarly, the homolog template 6RR7.1.A (3.01 Å) (https://www.rcsb.org/structure/6RR7)was used in order to build three dimensional model of PA protein.

Following the model preparation, these models were subjected to structure validation. The best model for each target protein among obtained models were estimated using parameters such as Ramachandran plot analysis using SAVES version 6 online tool(https://saves.mbi.ucla.edu/) and and z-score analysis through ProSA web server(https://prosa.services.came.sbg.ac.at/prosa.php).Autodock tools(Morris et al., 2009) was used for the protein preparation whereby hydrogen atoms were added, non-polar bonds being merged, Gasteiger charges being added and finally the proteins were converted to PDBQT format prior to docking.

### 2.3 Ligand library and ligand preparation

The FDA approved drugs, containing 1615 compounds were selected for the present study for virtual screening against HA and PA protein. The three dimensional structures of these FDA ligands were downloaded from the ZINC 15 database (https://zinc15.docking.org/) in Structural data file(SDF) format.(Sterling & Irwin, 2015). The ligands with favourable pharmacokinetic properties were filtered on the basis of Lipinski’s Rule of 5(Lipinski et al., 1997) and Veber’s rule using DATAWARRIOR software.(Sander et al., 2015). Openbabel GUI (O’Boyle et. al., 2011) available in PyRx 0.9.8 setup(Dallakyan & Olson, 2015) was used for the purpose of ligand preparation. The energy minimization of these compounds were performed and converted to PDBQT file format, a useable format for docking purpose. For the purpose of validating docking processes and parameters, native ligand of HA, oxamic acid has been used as a reference ligand. Similarly, 2-4-dioxo-4-phenylbutanoic acid, earlier identified as a potential drug candidate against PA protein has been used as a reference ligand. (Kowalinski et al. 2012)

### 2.4 hMAT1A filter for hepatotoxicity

hMAT1A is a gene expressed by mammals in normal liver. (Lu & Mato) hMAT1A filter is usually used to select the ligands from FDA library that won’t replace the native ligand SAM of HMAT1A. Hmat1A protein, Crystal structure of the human S-adenosylmethionine synthetase 1 (ligand-free form) (RCSB:6SW5)(https://www.rcsb.org/structure/6SW5) was converted to PDBQT format through Autodock tools before docking. Initial docking was performed between FDA ligand library and the hMAT1A protein. SAM(S-Adenosylmethionine) was taken as a reference inhibitor. The virtual screening was carried out whereby the ligands with binding energy lower than SAM inhibitor were taken and those with higher energy were filtered out.

### 2.5 Binding site prediction

The active binding pocket of the target protein HA was determined with the help of its homolog template 6D8W using Pymol software. Similarly,in case of PA,template 4AWF was used to identify the active binding residues.

### 2.6 Structure based Virtual Screening

The molecular docking was performed using Autodock Vina in PyRx (Dallakyan & Olson, 2015) with the above hMAT1A filtered ligands of FDA along with the reference/native ligand for each protein. The docking results were obtained as a negative value in Kcal/mol. The conformation that with the lowest docked energy are chosen as lowest docked energy corresponds to highest binding energy. The selection of top hit was done on the basis of highest preference index parameter and lowest docked energy for each target protein. The preference index, one of major parameter to prioritize drug candidates was calculated using formula in equation 1:

> PI=(Number of Hydrogen donor +Number of Hydrogen acceptor+Number of rotatable bonds)/25*5.

Following the molecular docking and identification of top hits for each HA and PA protein, the significant binding interactions between top leads and target protein were analysed using Chimera 1.15,Biovia Discovery Studio 2020 software (Baskaran et al. 2020) and LigPlus software(RA & MB, 2011)

### 2.7 Ligand-based ADME-Tox prediction

After identification of top hits for HA and PA target, the pharmacokinetic and druglikeness properties of these hits are further analysed. The online web tool SwissADME (http://www.swissadme.ch/index.php)(Daina et al., 2017) and pkCSM web tools(http://biosig.unimelb.edu.au/pkcsm/prediction) (Pires et al., 2015)were used to analyse various ADME as well as toxicological and pharmacokinetic properties properties of selected potential drugs.

### 2.8 Mutagenesis study

Mutagenesis study was performed to predict the probable mutation that is more likely to occur in the target protein. The probable mutations for HA protein were predicted and the effectivess of the identified top hits against mutated variants were analysed on the basis of comparision of wild binding energy with mutant binding energy. Using PyMoL, The predicted single and/or double mutation where single nucleotide was changed in single mutation and double nucleotides were changed in double mutation was performed using Pymol. It was based on the formation of rare tautomeric state Watson and Crick (WC) base pairs(Srivastava, 2019).

## 3. Results

### 3.1 Homology modeling

The strain of HA protein being selected for drug designing was: A/swine/Jiangsu/J006/2018(H1N1)–Uniprot accession number: MN416626. Similarly the strain A/swine/Jilin/23/2016(H1N1)has been selected to identify inhibitor against PA protein. (Brázda et al., 2021)

Due to the unavailability of the tertiary structure of HA and PA of G4 virus deposit in protein database, homology modeling was performed using Swiss model based on the target–template alignment. The parameters considered to identify the best models generated by Swiss model for each target protein is shown in **Table 1:**

**Table 1:**
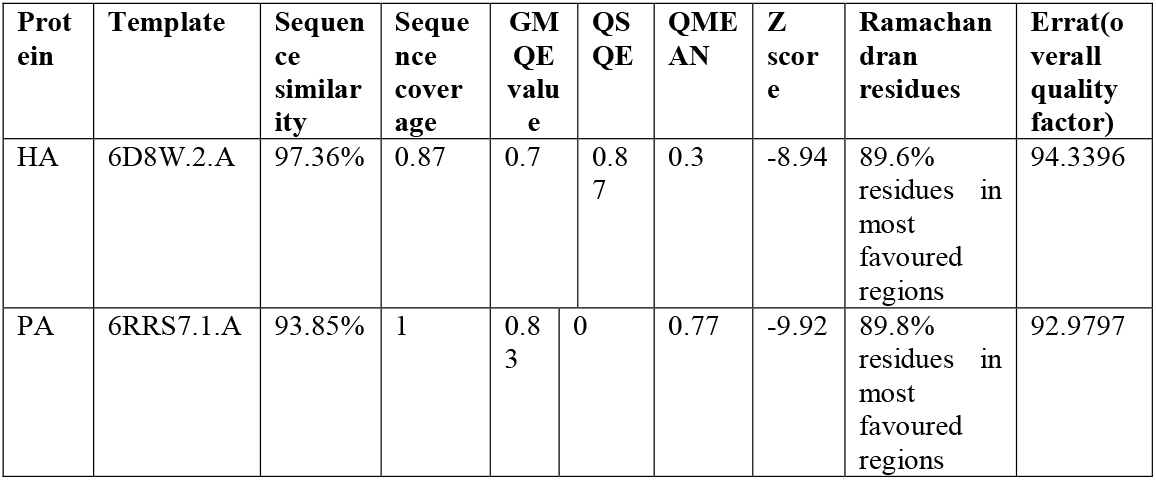
Homology modelling of HA and PA using Swissmodel and Structure validation.

### 3.2 Protein structure validation

The stability of the built models was analysed by the Structure analysis and verification server, SAVES v 6.0 (https://saves.mbi.ucla.edu/). The quality of each of the modeled protein was verified by the ERRAT score calculated in SAVES as shown in **Table 1** for each model. The overall stereochemical quality and overall residue by residue geometry of the built models were analysed by the PROCHECK, wherein Ramachandran plot for each model were analysed. Additionally, the z score of each model were calculated using ProSA web server(https://prosa.services.came.sbg.ac.at/prosa.php). The overall model quality is designated by the z score. The z score of each target proteins are shown in Table 1, where the energy of ProSA in negative value shows the reliability of the model, representing a good quality model.

### 3.3 hMAT1A filter of FDA Ligands

During identification of the best drug candidates from our FDA ligand library,these ligands are subjected to additional filter with HMAT1A protein using SAM as the inhibitor.Out of 192 filtered ligands of FDA ligands from Data warrior software,113 ligands were found to have binding energy less than that of SAM(native ligand of hMAT1A).The binding energy of SAM inhibitor was found to be −7.2 Kcal/mol. Thus, ligands with binding energy greater than that of SAM inhibitor, that could possibly replace SAM of HMAT1A has been filtered off and removed from ligand library prepared for molecular docking.

### 3.4 Molecular docking

#### 3.4.1 Identification of top hits

The structure based virtual screening performed between targets HA and PA with FDA approved compounds revealed several compounds interact very strongly with higher binding affinity. Out of 113 FDA ligands, the docking with HA protein revealed that all of those 113 ligands exhibited binding energy more than that of the reference/control ligand Oxamic acid(−4.2 Kcal/mole). The top 20 ligands on the basis of binding energy were taken for further analysis and hit identification. The preference index for each of top 20hits were also calculated based on inherent chemical property of the molecule as shown in **Supplementary Table 1 a.**. Based on the preference index, binding energy of the target with particular ligand and the binding energy with HMAT1A of these ligands, the top 3 hits were identified as presented in **Table 2 a**

**Table 2 a:** Top 3 hits of HA.

| Compound | Name | Preference Index | Binding energy against hmat1A(kcal/mol) | Binding energy against HA(kcal/mol) |
| --- | --- | --- | --- | --- |
| Compound_154 | Enalapril | 3.8 | -6.6 | -6.5 |
| Compound_22 | Enalaprilat | 3.6 | -6.8 | -6.5 |
| Compound_166 | Ivabradine | 3.4 | -6.4 | -6.7 |

Compounds 22 (Enalaprilat), 154(Enalapril) and 166(Ivabradine) showed high affinities towards the HA receptor, with docking scores of −6.5 Kcal/mol, −6.5 Kcal/mol and −6.7 KCal/mol, respectively, compared with a docking score of −4.2 kcal/mol for reference molecule Oxami cacid.

Similarly in the case of PA target, 113 Compounds were found to have binding energy higher than reference(−5.2 Kcal/mol). Top 20 hits were identified after docking as shown in **supplementary table 1(b).** On the basis of binding energy with ligands, HAMT1A protein with particular ligands and preference index, top 3 hits were identified as earlier. The hits identified are similar to that those identified for HA target as shown **in Table 2(b).**

**Table 2(b):**
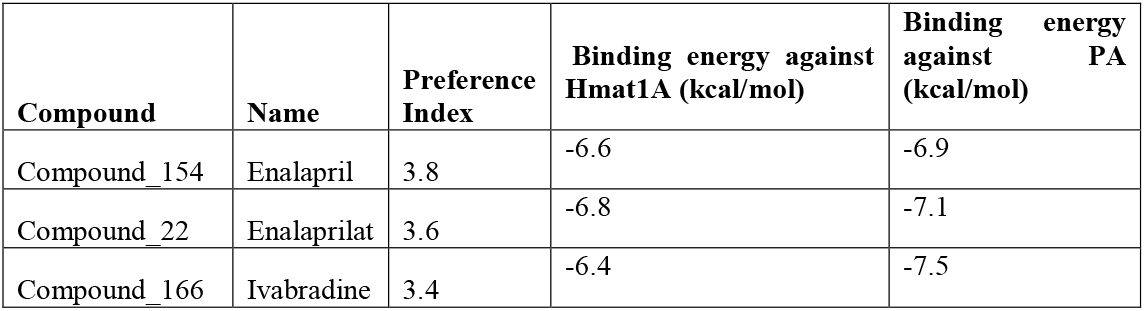
Top 3 hits of PA.

| Compound | Name | Preference Index | Binding energy against Hmat1A (kcal/mol) | Binding energy against PA (kcal/mol) |
| --- | --- | --- | --- | --- |
| Compound_154 | Enalapril | 3.8 | -6.6 | -6.9 |
| Compound_22 | Enalaprilat | 3.6 | -6.8 | -7.1 |
| Compound_166 | Ivabradine | 3.4 | -6.4 | -7.5 |

The docking result revealed that the Compounds 22 (Enalaprilat), 154(Enalapril) and 166(Ivabradine) showed high affinities towards the PA receptor as well, with docking scores of − 7.1 Kcal/mol,−6.9 Kcal/mol and −7.5 KCal/mol, respectively, compared with a docking score of - 5.2kcal/mol for its reference molecule.

#### 3.4.2 All respective top hits were found to bound the binding groove of HA and PA protein

Following the molecular docking and identification of top hits, the interactions exhibited by the drug and the targets were visualized using Chimera 1.15 software and Biovia Discovery Studio. The respective identified drugs were found to be bound at the desired binding groove of HA and PA protein as revealed by the Pymol software. Moreover, the interacting residues of the target protein HA and PA with the respective top 3 putative drug candidates were analyzed as shown **in Supplementary Table 2 a**.Here, the hydrogen, carbon –hydrogen bond and other interactions were analayzed using Discovery Studio and hydrophobic bonds being analysed using LipPlot+.

### 3.5 MUTATION ANALYSIS

The occurance of spontaneous point mutation, transitions of the base pairs G:C to A:T and A:T to G:C, are inherent characteristics of DNA.(Trixler, 2013).The constant point mutations could result in the mutations in the antigenic sites and lead to the evolution of viruses which ultimately produce new viral subtypes(Carrat & Flahault, 2007). The nucleic acid bases follow the Watson-Crick (WC) pairing rule. But however due to changed tautomeric and anionic form, the usual Watson –Crick pairing can be altered. The probable spontaneous point mutation most likely to occur were predicted by changing the first or second nucleotides in singe mutation and changing both nucleotides of a codon in double mutation as shown in **Supplementary Table 3a.**

The Alanine at the position 241 of HA is predicted to mutate to either to Threonine or Valine or Isoleucine. Similarly, the Glycine at position 242 is predicted to undergo mutation, resulting in either Serine or Aspartic acid or Asparagine at this particular position. Besides, the 159Asparagine forming conventional hydrogen bond with ligand 22 and ligand 154, might undergo mutation as well. However, the change in either of the 3 nucleotides of the codon coding for 159Asparagine still results in generation of Asparagine itself due to Wobble phenomenon of codons (Yarus, 2021).Additionally, the mutation in 240Glutamine,hydrogen bond with the ligand, results in the generation of Stop Codon (TAA), resulting in truncated protein which will render the protein defunct and could be lethal to virus. Such derived mutant proteins were further subjected to molecular docking with the top hits to examine the ligands binding affinity to the respective mutants and whether these narrowed ligands could also be effective towards the respective variant virus.

**Table 3a:** Molecular docking of HA mutated protein structures with top hits, showing wild binding energy and mutant binding energy.

| **Protein;Residue** | **Mutation** | **Ligands** | **Mutant B.E** | **Wild B.E.** | **P.I** |
| --- | --- | --- | --- | --- | --- |
| Alanine241 | A241I | compound_166 | -6.7 | -6.7 | 3.4 |
|  |  | compound_22 | -6.4 | -6.5 | 3.6 |
|  |  | compound_154 | -6.3 | -6.5 | 3.8 |
|  |  | oxamic acid 974 | -4.2 | -4.2 |  |
| Alanine241 | A241T | compound_166 | -6.7 | -6.7 | 3.4 |
|  |  | compound_154 | -6.6 | -6.5 | 3.8 |
|  |  | compound_22 | -6.5 | -6.5 | 3.6 |
|  |  | oxamic acid 974 | -4.2 | -4.2 |  |
| Alanine241 | A241V | compound_166 | -6.6 | -6.7 | 3.4 |
|  |  | compound_22 | -6.4 | -6.5 | 3.6 |
|  |  | compound_154 | -6.3 | -6.5 | 3.8 |
|  |  | oxamic acid 974 | -4.2 | -4.2 |  |
| Glycine242 | G242S | compound_22 | -6.5 | -6.5 | 3.6 |
|  |  | compound_154 | -6.2 | -6.5 | 3.8 |
|  |  | compound_166 | -6.2 | -6.7 | 3.4 |
|  |  | oxamic acid 974 | -4.2 | -4.2 |  |
| Glycine242 | G242D | compound_166 | -6.7 | -6.7 | 3.4 |
|  |  | compound_154 | -5.9 | -6.5 | 3.8 |
|  |  | compound_22 | -5.9 | -6.5 | 3.6 |
|  |  | oxamic acid 974 | -4.2 | -4.2 |  |
| Glycine242 | G242N | compound_22 | -6.2 | -6.5 | 3.6 |
|  |  | compound_154 | -5.9 | -6.5 | 3.8 |
|  |  | compound_166 | -5.7 | -6.7 | 3.4 |
|  |  | oxamic acid 974 | -4.2 | -4.2 |  |

The binding energies of the top hit ligands to the wild type and the protein with residue at which the mutation has been predicted at the respective codons after docking are presented in Table 4a. The calculated preference indices (PI) for each of the top 3 hits are presented as well. The docking analysis of each mutated protein revealed all the top three leads for HA protein work effectively for all the respective mutant proteins, too. The binding energy for each mutated protein was found to be greater than the binding energy of the reference ligand Oxamic acid. The Enalapril, Enalaprilat and Ivabradine were effective for the HA protein even after undergoing probable mutations suggesting that the drug could be effective even for these probable variants.

Moreover, among these ligands compound 154, Enalapril, had the highest preference index indicating of better pharmacokinetic properties of the drug in compared to rest of the hits. Hence, the compound 154, Enalapril, has been identified as the best hits among the FDA ligands for wild as well as predicted mutant HA proteins.

**Table 3b:**
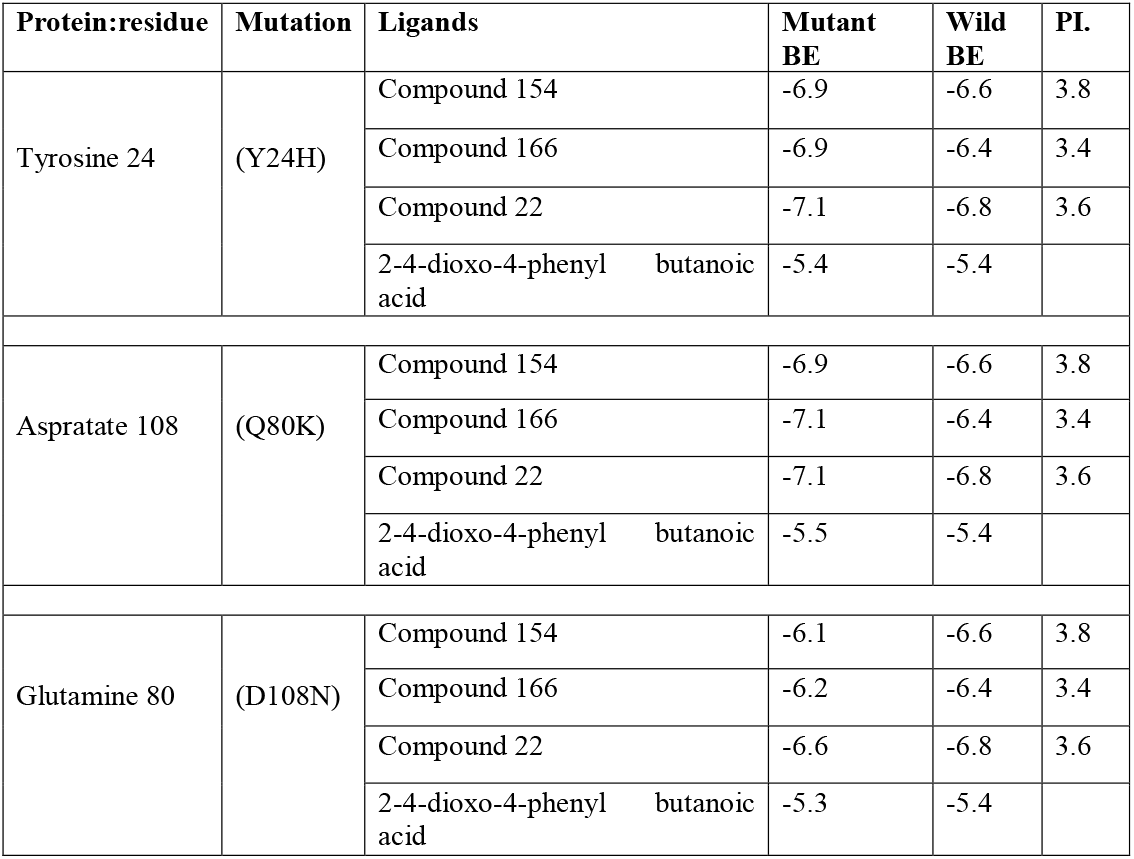
Molecular docking of PA mutated protein structures with top hits, showing wild binding energy and mutant binding energy.

All the mutated protein were found to have higher binding energy than the reference ligand 2-4 dioxo-4phenyl butanoic acid (−5.4 k/mol) and the top hits (Enalapril, Enalaprilat and Ivabradine) could be effective against these mutated protein as well. Thus, it can be inferred from the above table that the preference index of compound 154, Enalapril, is higher than the other compound and has higher binding energy than the reference ligand. Compound154, Enalapril has been selected as the best hits for the wild and mutated protein of PA.

These results clearly demonstrate that Enalapril is could be drug candidate as it can act against HA and PA along with the predicted respective mutants of active sites of the respective proteins. Thus, Enalapril could be a drug candidate to treat EA H1N1 Swine Flu G4 variant as this targets multiple important proteins critical for viral survival, replication and infectivity.

## 4. Discussion

The emerging global threat of influenza viruses is the result of continuous evolution due to various reasons. Mutation (antigenic drift), reassortment (antigenic shift) and recombination are the major mechanism responsible for the evolution in the influenza virus(Shao et al., 2017).The formation of new subtypes of influenza virus plays a significant role in wide propagation and disastrous pandemics as there is dearth of immunity to the newly emerging subtypes(Ma et al., 2008).It is prudent that the occurrence and subsequent outbreak of such lethal pathogenic viruses is a major global health threat. The vaccination has been used as a staple method against epidemic and pandemic influenza. However, it is not able to provide the immediate protection against such epidemic and pandemic flu outbreaks. The development of antiviral agents with immunomodulatory characteristics as a supplement to vaccination is very crucial to combat the pandemics(Shao et al., 2017). On another note, *in silico* modelling and computational biology approach could be an inevitable approach in identifying lead potential Nobel drugs for the alleviation of particular disease. Drug repurposing, an strategy for identifying new uses for approved or investigational drugs could be employed. For drug discovery process, the protein target selection is pivotal. HA and PA proteins have been selected as a target proteins against whose putative drug candidates were sought.

Haemagglutinin is a protein responsible for viral entry to the host, thus, determining pathogenicity and mediating membrane fusion.It has a great role in antigenic drift and shift as well and is assumed to be an important target for drug discovery(Wu & Wilson, 2020). The HA protein has now already been established as a validated drug target against influenza virus. However, no any FDA-approved therapeutics have been succeeded in specifically blocking the receptor binding or the fusion machinery to prevent the functions of HA(Yao et al., 2020). PA is RNA dependent RNA polymerase of influenza virus that functions as endonuclease. (Bhoye and Cherian 2018).It cleaves the host mRNAs downstream of their mRNA cap structures which are recognized and bound by PB2. (S. Yuan et al. 2015)

Homology modeling was employed to prepare the three dimensional structure of the respective target proteins using SwissModel online tool. The best quality model was identified using GMQE value,that gives an overall model quality measurement and lies between 0 and 1,where higher numbers indicates the higher expected quality of model. Further, the QMEAN (Qualitative Model Energy Analysis) score of the built models were considered that is a composite scoring function describing the major geometrical aspects of built protein structures (P et al., 2008).The QMEAN structure of the generated model was found to be less than 1 for all proteins, indicating these built models are comparable to their respective template structures(Aatif et al., 2021).The stability of the built model was analysed by the Structure analysis and verification server, SAVES v 6.0 (https://saves.mbi.ucla.edu/). The quality of each of the modeled protein was verified by the ERRAT score calculated in SAVES, where the value depicted that the non-bonded interactions of each models lied within a reasonable normal range(Barge et al., 2021)The overall stereochemical quality and overall and residue by residue geometry of the build models are analysed by the Procheck, wherein the Ramachandran plot were analysed for each models. The Ramachandran plot is basically used to check the protein conformation and reliability of φ and ψ angles in the backbone of amino acid residues in protein(Barge et al., 2021; Rakib et al., 2021)

Following protein structure preparation, prior to molecular docking with aim to identify the best hit, hMAT1A filter was employed with reference S-Adenosyl Methioine (SAM). native ligand of HMAT1A). MAT1A is a gene expressed by mammals in normal liver.The MAT1A gene encodes a enzyme methionine adenosyl transferase. Methionine adenosyl transferase (MAT) protein is the enzyme that catalyzes the biosynthesis of AdoMet(SAM) from ATP and methionine, usually express by the mammal in normal liver.(Lu & Mato, 2012) Only ligands with binding energy less than SAM inhibitor were taken such that our putative drug candidates won’t replace the native ligand SAM of HMAT1A that would otherwise, disrupts cellular functions and causes hepatotoxicity.

Molecular docking is one of the tools to screen for the ligands that could potentially act as competitive inhibitor as a drug candidate against the target. Out of 113 FDA ligands filtered from Hmat1A docking, the docking with HA protein revealed that all of those 113 ligands exhibited binding energy more than that of the reference/control ligand Oxamicacid(−4.2 Kcal/mole).The top 20 ligands were taken for further analysis and hit identification based on binding energy. Similarly, 107 compounds of PA was found to have higher binding energy than its reference. The top 20 compounds were identified and sorted as in **Supplementary Table 1 b** on the basis of preference index, Hmat 1 A binding energy with that ligand and its binding energy with respective target. The top 3 compounds were selected on the basis of these 3 parameters along with consideration of their respective pharmacokinetic properties.

To identify potential medication candidates, PI has been employed as a key metric. Hydrogen bond, hydrophobic bond, and rotable bond count are all taken into account in equation 1.The binding affinity between the drug and the target proteins is increased by hydrophobic bonds.The formation of a hydrogen bond is necessary for the protein-ligand combination to be stable. Furthermore, drug selectivity, metabolization, and adsorption in the body are all governed by hydrogen bonding.(Barge et al., 2021) The total polar surface area, which aids in membrane permeability, is another element of the preference index.

Following the molecular docking and identification of top hits, the interactions exhibited by the drug and the targets were visualized using Pymol software. The pymol software revealed that the identified medicines were bound to the desired binding groove of the HA and PA proteins. The drugs identified for HA targets the receptor binding domain and N-terminal domain of PA protein. We can deduce from these findings that medications would inhibit these two proteins.

The inherent property of the influenza viruses to undergo spontaneous mutation could be explained by the phenomenon known as quantum tunneling, more specifically, proton tunneling.In accordance with the wave –particle dual nature of particular particle, in the sub atomic level the electrons behave like wave like particles such that electrons tunnel through an obstacle rather than climbing over it(*Schrôdinger’s Mutations – TheGIST*, n.d.) Proton tunneling has been reported for the spontaneous mutation inside the genome by various simulation methods and quantum approaches. Here, the proton disappears from one position and reappears nearby simultaneously. The hydrogen bonds between two strands of DNA helix, under certain conditions behave like waves and spread to many locations at once by proton tunneling phenomena. This is what is thought to be leading to spontaneous mutation due to occasional finding of these atoms on wrong strand of DNA (A new study reveals that quantum physics can cause mutations in our DNA | University of Surrey n.d.,(Löwdin, 1963)). Moreover, the subsequent errors during replication could be predicted by the basic knowledge of various quantum jump events. (Mcfadden & Al-Khalili, 1999; Strippoli et al., 2005)

Thus, it is assumed that probable mutant prediction and drug screening based on affinity towards wild type and possible mutant proteins with the drugs preference index could be taken as one of the parameters to prioritize the drug candidate in treating disease for which effective medication might not have been yet discovered or prescribed.

## 5. Conclusions

It can be suggested that screening of hit ligand with higher binding energy towards wild type and predicted active site amino acids mutant proteins with higher preference index (pharmacophore properties and probable reactivity of the ligand towards the amino acids) could be screened using CADD approach of molecular docking and probable mutant prediction as template molecule to treat disease or symptom management for which an effective drug has not yet been discovered or prescribed.

---

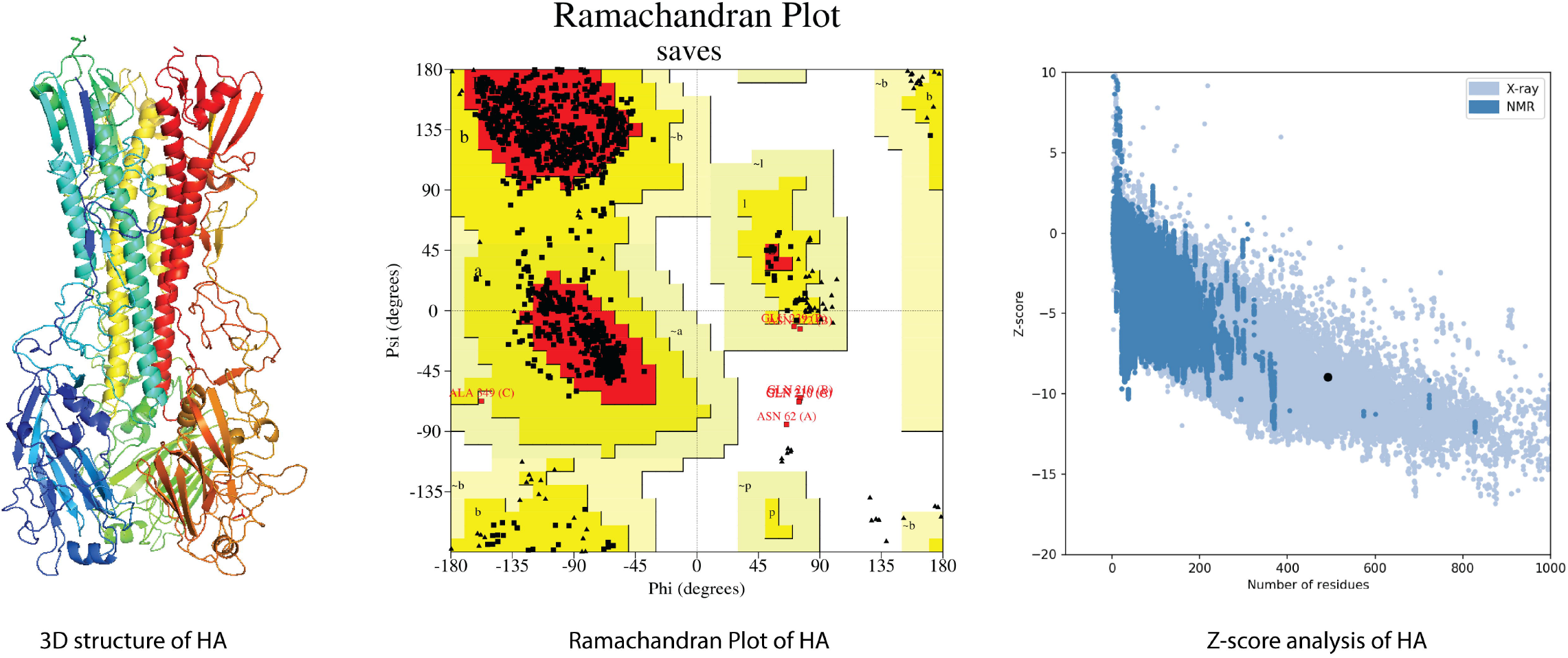

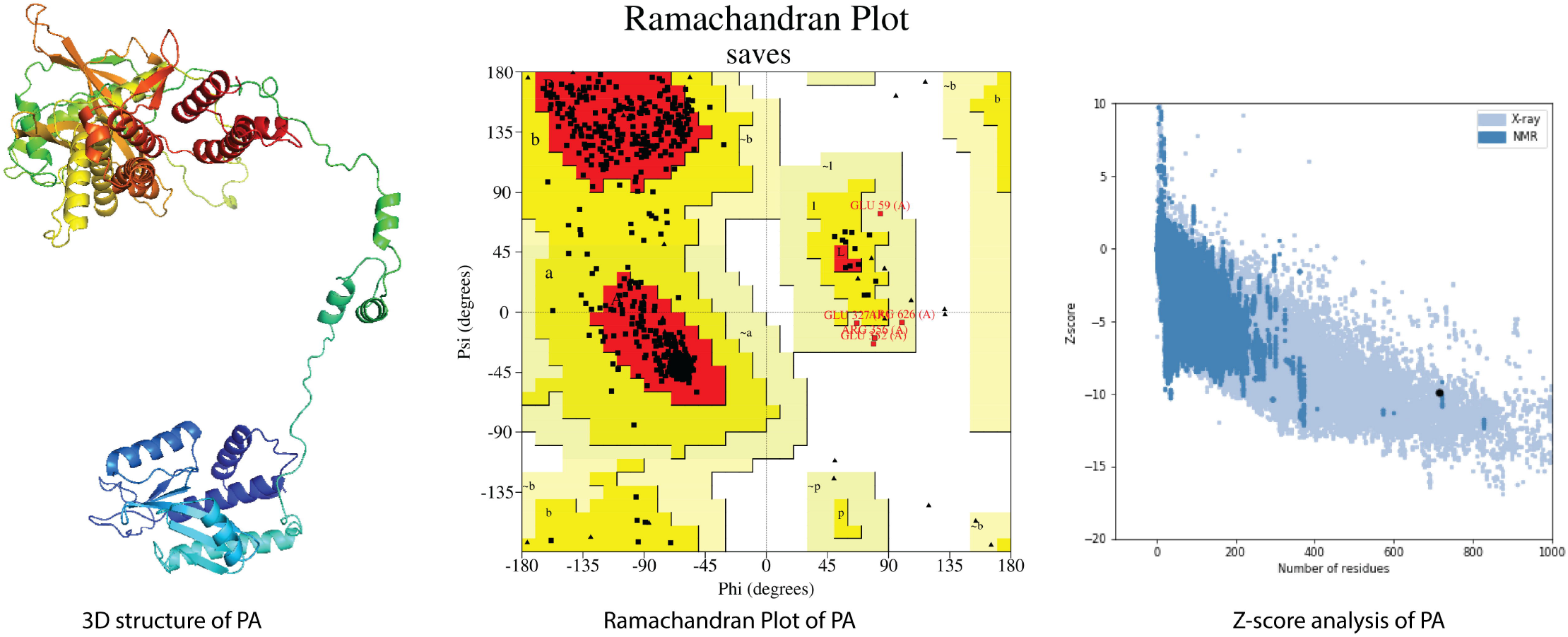

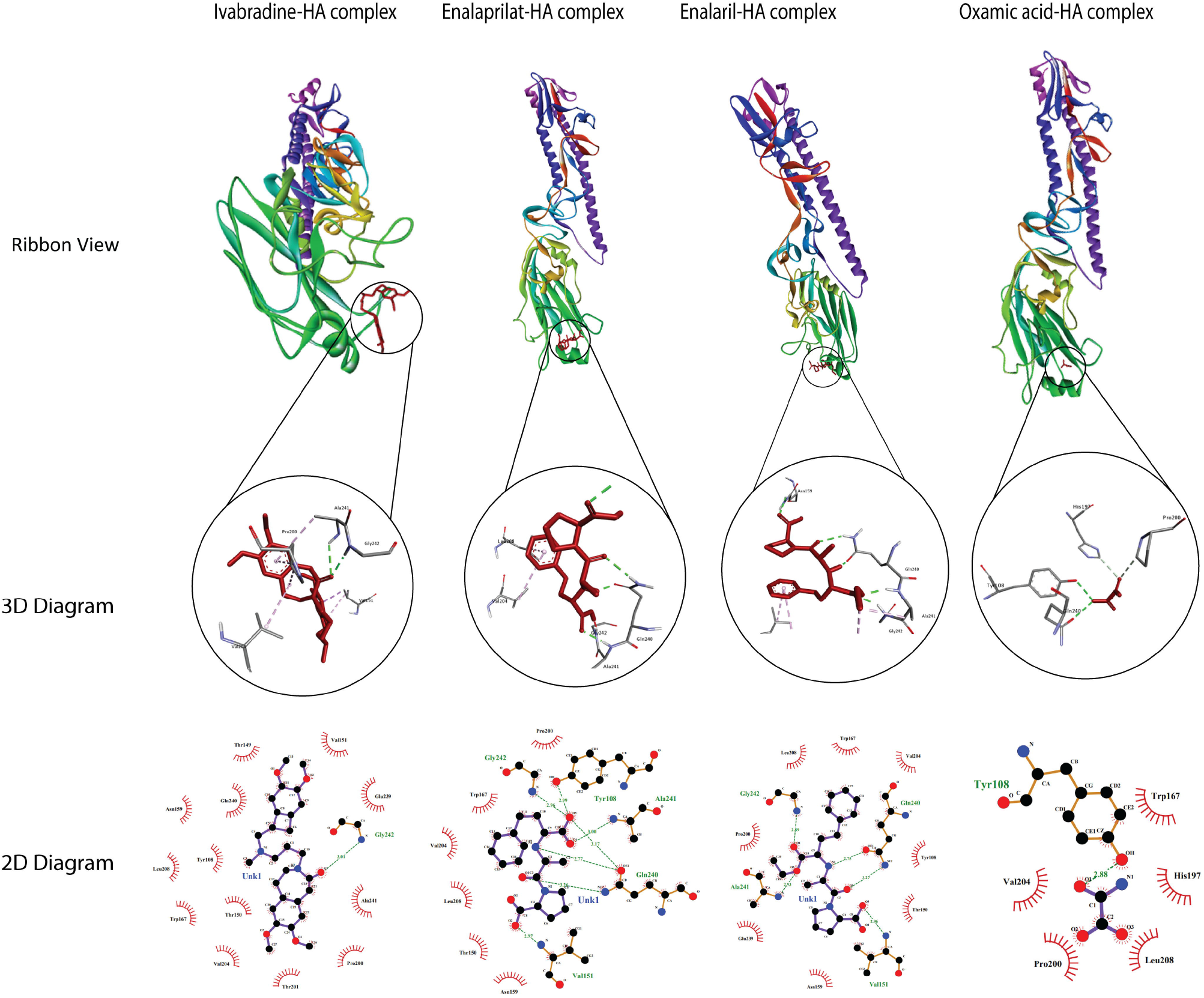

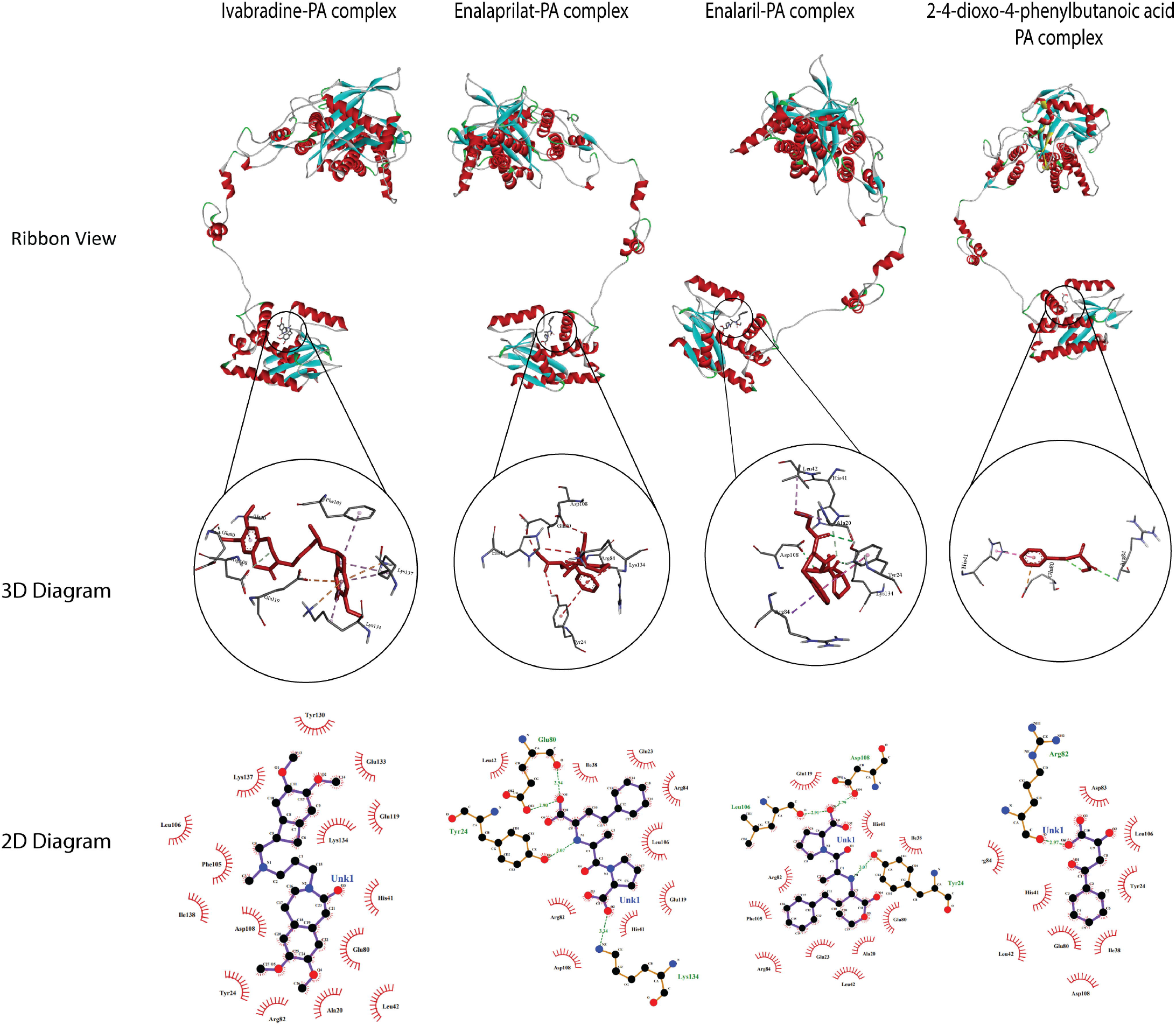

