## Supplemental file for "Repurposing of Drugs Against Mutated Strain of Eurasian Avian Like H1N1 (EA H1N1) Swine Flu Virus, Genotype 4(G4) Virus"

**Supplementary**

**Supplementary Table 1 a) Top 20 hits of HA protein along with its parameters considered**

| **Compound** | **Name** | **Hydrogen Acceptor** | **Hydrogen Donor** | **Rotable bonds** | **Preference Index** | **Binding energy with Hmat1A** | **Binding energy against HA** |
| --- | --- | --- | --- | --- | --- | --- | --- |
| Compound_154 | Enalapril | *7* | *2* | *10* | *3.8* | *-6.6* | *-6.5* |
| Compound_17 | Alfuzosin | 9 | 2 | 8 | 3.8 | -7 | -5.8 |
| Compound_189 | (S,R) Formoterol | 6 | 4 | 8 | 3.6 | -7.2 | -7.2 |
| Compound_106 | (S,S)-Formoterol | 6 | 4 | 8 | 3.6 | -7.2 | -7.1 |
| Compound_10 | (R,R)-Formoterol | 6 | 4 | 8 | 3.6 | -7 | -7 |
| Compound_188 | (R,S)-Formoterol | 6 | 4 | 8 | 3.6 | -7.2 | -6.8 |
| Compound_22 | *Enalaprilat* | *7* | *3* | *8* | *3.6* | *-6.8* | *-6.5* |
| Compound_166 | *Ivabradine* | *7* | *0* | *10* | *3.4* | *-6.4* | *-6.7* |
| Compound_37 | (S)-Rosiglitazone | 6 | 1 | 7 | 2.8 | -6.4 | -6.4 |
| Compound_43 | Voriconazole | 6 | 1 | 5 | 2.4 | -6.7 | -6.6 |
| Compound_191 | Acrivastine | 4 | 1 | 6 | 2.2 | -7 | -6.7 |
| Compound_125 | ZINC100036756 | 5 | 1 | 5 | 2.2 | -7 | -6.7 |
| Compound_48 | ZINC100296828 | 4 | 1 | 6 | 2.2 | -7.1 | -6.7 |
| Compound_19 | Tiotropium | 5 | 1 | 5 | 2.2 | -7.2 | -6.4 |
| Compound_143 | Scopolamine | 5 | 1 | 5 | 2.2 | -7.2 | -6.1 |
| Compound_121 | Ipratropium | 4 | 1 | 6 | 2.2 | -7 | -5.8 |
| Compound_126 | ZINC100027531 | 5 | 1 | 5 | 2.2 | -7.1 | -5.8 |
| Compound_7 | L-Naloxone | 5 | 2 | 2 | 1.8 | -7.1 | -6.5 |
| Compound _4 | Frovatriptan | 4 | 3 | 2 | 1.8 | -7.1 | -6.2 |
| Compound _38 | Hydrocodone | 4 | 0 | 1 | 1 | -7 | -6.6 |

**Supplementary Table 1 b) Top 20 hits of PA protein along with its parameters considered**

| **Compound** | **Name** | **Hydrogen Acceptor** | **Hydrogen Donor** | **Rotable bonds** | **Preference Index** | **Hmat1A Binding energy** | **Binding energy** |
| --- | --- | --- | --- | --- | --- | --- | --- |
| Compound_154 | Enalapril | *7* | *2* | *10* | *3.8* | *-6.6* | *-6.9* |
| Compound_17 | Alfuzosin | 9 | 2 | 8 | 3.8 | -7 | -6.6 |
| Compound_22 | Enalaprilat | *7* | *3* | *8* | *3.6* | *-6.8* | *-7.1* |
| Compound_166 | Ivabradine | *7* | *0* | *10* | *3.4* | *-6.4* | *-7.5* |
| Compound_99 | Zymar | 7 | 2 | 4 | 2.6 | -7 | -6.7 |
| Compound_43 | voriconazole | 6 | 1 | 5 | 2.4 | -6.7 | -6.5 |
| Compound_48 | ZINC000100296828 | 4 | 1 | 6 | 2.2 | -7.1 | -8 |
| Compound_187 | Efinaconazole | 5 | 1 | 5 | 2.2 | -6.8 | -6.8 |
| Compound_191 | Acrivastine | 4 | 1 | 6 | 2.2 | -7 | -7.8 |
| Compound_19 | Tiotropium | 5 | 1 | 5 | 2.2 | -7.2 | -6.4 |
| Compound_126 | Pamine | 5 | 1 | 5 | 2.2 | -7.1 | -6.5 |
| Compound_143 | scopolamine | 5 | 1 | 5 | 2.2 | -7.2 | -6.3 |
| Compound_15 | Rozerem | 3 | 1 | 4 | 1.6 | -7 | -7 |
| Compound_162 | Ketorolac | 4 | 1 | 3 | 1.6 | -7.1 | -6.8 |
| Compound_45 | Ketorolac | 4 | 1 | 3 | 1.6 | -7.1 | -6.5 |
| Compound_137 | Brimonidine | 5 | 2 | 1 | 1.6 | -6.6 | -6.4 |
| Compound_52 | Triprolidine | 2 | 0 | 4 | 1.2 | -7 | -7.6 |
| Compound_90 | Galantamine | 4 | 1 | 1 | 1.2 | -7 | -6.7 |
| Compound_164 | Anagrelide | 4 | 1 | 0 | 1 | -6.9 | -6.5 |
| Compound _28 | Esmirtazapine | 3 | 0 | 0 | 0.6 | -7.1 | -7.7 |

**Supplementary Table 2a: Interacting residues of HA target with top 3 hits**

| Compound | Interacting Residues | Hydrogen Bond  Residues | Hydrophobic Bond  Residues | B.E  Kcal/mol |
| --- | --- | --- | --- | --- |
| 154  Enalapril | Tyr 108, Asp144, Arg 147,Gly148,  Thr 149, Thr 150, Val 151,Ala152,  Asn 159, Trp 167, Val 169,Lys 170,  Gly 172, His 197, Pro 199, Pro200,  Thr 201, Ser 203, Val204, Thr207,  Leu 208,Lys 236,Glu 239,Gln 240,  Ala 241,Gly 242 | Ala 241,  Gly 242,  Gln 240,  Asn 159 | Leu208, Trp 167,  Val 204, Pro 200,  Glu 239,Asn 159,  Thr 150, Tyr 108,  Gly 242, Ala 241,  Gln 240, Val 151 | -6.5 |
| 22  Enalaprilat | Tyr108, Asp144, Arg147, Gly148,  Thr149, Thr150, Val151, Ala152,  Asn159,Trp 167,Val 169, Lys 170,  Gly172, His197, His198, Pro 199,  Pro200,Thr 201,Ser 203,Val 204,  Thr 207,Leu 208,Lys 236,Glu 239,  Gln 240,Ala 241,Gly 242,Arg 243 | Ala 241,  Asn 159  Gln 240 | Leu 208, Val 204,  Trp 167, Pro 200,  Asn 159, Thr 150,  Gly 242, Tyr 108,  Gln 240, Val 151 | -6.5 |
| 166  Ivabradine | Tyr 108,Asp144,Arg 147,Gly 148,  Thr 149,Thr 150,Val 151,Ala 152,  Asn 159,Trp 167,Val 169,Lys 170,  Gly 172,Lys 177,His 197,Thr 201,  Ser 203,Val 204,Thr 207,Leu 208,  Ala 233,Lys 236,Glu 239,Gln 240,  Ala 241,Gly 242 | Gly 242,  Ala 241 | Thr149, Val151,  Gln240, Asn159,  Tyr 108, Leu 208,  Trp167, Val204,  Thr201,Pro 200,  Ala 241,Glu 239,  Thr 150,Gly 242 | -6.7 |
| Oxamic acid  Reference | Tyr 108,Thr 150,Trp 167,His 197,  Pro 199,Pro 200,Val 204,Leu 208,  Gln 240,Gly 242 | Tyr 108,  Gln 240 | Leu208, Pro 200,  Val204, His 197,  Trp 167, Tyr 108 | -4.2 |

**Supplementary Table 2 b : Interacting residues of PA target with top 3 hits**

| Compound | Interacting Residues | Hydrogen Bond  Residues | Hydrophobic Bond  Residues | B.E  Kcal/mol |
| --- | --- | --- | --- | --- |
| 154  Enalapril | Glu133,Lys137,Lys104,Ile138,  Lys134,Tyr130,Phe105,Leu106,  Pro107,Glu119,Trp88,Ala87,  Ile79,Gly121,Arg84,Ile120  Asp108,Asp83,Glu80,Arg82,  Gly81,Try24,Glu23,Val122,  Ala20, His41,Leu42,Leu16,  Ala17,Lys19,Ile38,Ala37,Lys34 | Lys134,  Tyr24,  Asp108 | Glu119, His41,  Ile38, Glu80,  Ala20 ,Leu42,  Glu23, Arg84,  Phe105,Arg82 | -6.6 |
| 22  Enalaprilat | Lys104,Ile138,Lys137,Phe105,  Lys134,Tyr130,Trp88,Leu106,  Pro107,Glu119,Ile85,Ala87,  Arg84, Asp108, Ile79,Ile120,  Asp83,Glu80, Arg82,Gly81,  Tyr130,Tyr24,Gly121,Glu23,  Ala20,Val122,His41,Leu42,  Leu16,Ala17,Ile38,Ala37,Lys34 | Tyr24,  Glu80,  Asp108,  Lys134 | Leu42,Ile38,  Glu23,Arg84,  Leu106,Arg82,  Asp108,His41,  Glu119 | -6.8 |
| 166  Ivabradine | Lys34,Val122,Thr123,Tyr130,  Glu133,Lys134,Lys137,Ile138,  Ile38, His41,Glu23,Ala20,  Leu42, Ala17, Leu16, Gly121,  le120,Gly81, Arg82, Asp83.  Glu80, Ile79, Ala87, Arg84,  Asp108,Glu119,Pro107,Trp88,  Leu106,Phe105,Lys104,Tyr24 | Nil | Tyr130,Glu133,  Lys137,Glu199,  Lys134,Leu106,  Phe105,Ile138,  Asp108,His41,  Glu80,Leu42,  Ala20,Arg82,  Tyr24 | **-**6.4 |
| 2-4-dioxo-4-phenylbutanoic acid | Lys134,Phe105,Leu106,Glu119,  Pro107,Trp80,Ile120,Arg84,  Asp108, Ile79,Ala87,Glu80,  Tyr24, Glu23, Ile38, Ala20,  His41, Asp83,Arg82, Gly81,  Leu42, Ala17,Leu16 | Arg84 | Asp83,Leu106,  Tyr24, Ile38,  Asp108,Glu80,  Leu42,His41,  Arg84 | -5.4 |

**Supplementary Table 3a:Predicted mutation of HA protein**

| **Residue;preferred codons** | **Converted residue;Predicted mutation** | **Codons for predicted mutation** |
| --- | --- | --- |
| Alanine 241( GCA) | Threonine (A241T) | ACA (first nucleotide of the code) |
|  | Valine (A241V) | GTA (second nucleotide of the code) |
|  | Isoleucine (A241I) | ATA (first and second nucleotides of the code) |
| Glycine 242(GGC) | Serine (G242S) | AGC(first nucleotide of the code), AGT(first and third nucleotide of the code) |
|  | Aspartic Acid (G242D) | GAC(first nucleotide of the code) , GAT( second and third nucleotide of code) |
|  | Asparagine (G242N) | AAC( first and second nucleotide of code), AAT (all nucleotide of code) |

**Supplementary Table 3b :Predicted mutation of PA protein**

| **Residue;preferred codons** | **Converted residue;Predicted mutation** | **Codons for predicted mutation** |
| --- | --- | --- |
| Tyrosine 24(TAT) | Histidine (Y24H) | CAT (first nucleotide of code) |
| Glutamine 80(GAA) | Lysine (Q80K) | AAA (first nucleotide of code) |
| Aspartic Acid 108(GAT) | Asparagine (D108N) | AAT (first nucleotide of code) |
